# Minimal structural elements required for midline repulsive signaling and regulation of *Drosophila* Robo1

**DOI:** 10.1101/2020.03.30.016766

**Authors:** Haley E. Brown, Timothy A. Evans

## Abstract

The Roundabout (Robo) family of axon guidance receptors has a conserved ectodomain arrangement of five immunoglobulin-like (Ig) domains plus three fibronectin (Fn) repeats. Based on the strong evolutionary conservation of this domain structure among Robo receptors, as well as *in vitro* structural and domain-domain interaction studies of Robo family members, this ectodomain arrangement is predicted to be important for Robo receptor signaling in response to Slit ligands. Here, we define the minimal ectodomain structure required for Slit binding and midline repulsive signaling *in vivo* by *Drosophila* Robo1. We find that the majority of the Robo1 ectodomain is dispensable for both Slit binding and repulsive signaling. We show that a significant level of midline repulsive signaling activity is retained when all Robo1 ectodomain elements apart from Ig1 are deleted, and that the combination of Ig1 plus one additional ectodomain element (Ig2, Ig5, or Fn3) is sufficient to restore midline repulsion to wild type levels. Further, we find that deleting four out of five Robo1 Ig domains (ΔIg2-5) does not affect negative regulation of Robo1 by Commissureless (Comm) or Robo2, while variants lacking all three fibronectin repeats (ΔFn1-3 and ΔIg2-Fn3) are insensitive to regulation by both Comm and Robo2, signifying a novel regulatory role for Robo1’s Fn repeats. Our results provide an *in vivo* perspective on the importance of the conserved 5+3 ectodomain structure of Robo receptors, and suggest that specific biochemical properties and/or ectodomain structural conformations observed *in vitro* for domains other than Ig1 may have limited significance for *in vivo* signaling in the context of midline repulsion.

## Introduction

During development of the central nervous system (CNS), evolutionarily conserved ligands and receptors guide growing axons toward their synaptic targets via local regulation of actin dynamics in the axonal growth cone. Although the four major canonical axon guidance pathways (Ephrin-Eph, Netrin-DCC, Slit-Robo, Sema-Plexin) have been known since the 1990s [1,2], mechanistic details of ligand-receptor interaction, receptor signaling, and receptor regulation are still emerging [3,4].

Roundabout (Robo) family receptors and their cognate Slit ligands represent one of the best characterized repulsive axon guidance signaling pathways and are conserved across bilaterians, including insects, nematodes, planarians and vertebrates [5-10]. Most Robo family members, including the three *Drosophila* Robos (Robo1, Robo2 and Robo3), share a conserved 5+3 protein structure which consists of five Immunoglobulin-like (Ig) domains, three Fibronectin (Fn) type-III repeats, a transmembrane domain and two to four conserved cytoplasmic motifs (CC0, CC1, CC2, and CC3) [7,11]. With the exception of vertebrate Robo3/Rig-1 (and possibly Robo4/Magic Roundabout) [12-15], all Robo family members bind Slit ligands with high affinity. Previous studies have pinpointed this binding interface to the Ig1 domain of Robo and the second leucine-rich repeat (LRR D2) of Slit [6,16-20], and *in vivo* studies have confirmed the importance of Robo1 Ig1 for midline repulsive signaling [21,22].

Slit-Robo signaling *in vivo* is thought to involve a series of events including Slit binding, receptor cleavage and endocytosis, and recruitment of downstream effectors, although the mechanism(s) of receptor activation are not yet fully understood [11,23-26]. Robo receptors share structural similarity with cell adhesion molecules and lack intracellular catalytic domains [27]. One outstanding question is what relationship the characteristic Robo ectodomain structure (5 Ig + 3 Fn) has to Slit-dependent signaling mechanism(s). Homophilic and heterophilic interactions between Robo receptors have been reported [28-30], and *in vitro* evidence suggests that Robo ectodomains can adopt multimeric configurations in solution as well as at the cell surface in the absence of Slit [31-33], and that the dimerization state of Robo is unaltered by Slit binding [31,34]. The *in vivo* significance of these structural conformations and domain-domain interactions for receptor signaling and regulation remains unclear.

*Drosophila* Robo1’s axon guidance role has been best characterized in the embryonic ventral nerve cord, where it is responsible for preventing axons from inappropriately crossing the midline to the opposite side of the body. When Robo repulsion is lost, axons inappropriately cross and recross the midline forming characteristic circular roundabouts [7,35]. As many commissural axons need to cross the midline once to innervate the contralateral side of the body, precise temporal regulation of Slit-Robo repulsion is essential. Early in development *Drosophila* Robo1 is expressed only at low levels on the surface of pre-crossing commissural axons due to Commissureless (Comm) activity. This protein endosomally sorts Robo1 into Comm-containing vesicles to be targeted for degradation by the lysosome [36-42]. Even when *comm* is active, some Robo1 is able to escape degradation and be trafficked to the growth cone surface. To ensure axons can still cross the midline Robo1 is then further inhibited via interactions with the first and second Ig domains of Robo2 expressed by midline cells [29]. It has yet to be discovered where on Robo1 this trans-interaction takes place. After midline crossing, *comm* expression is terminated and Robo1 levels increase on growth cones of post-crossing commissural axons, preventing them from re-crossing.

The strong evolutionary conservation of ectodomain structural arrangements combined with *in vitro* evidence for domain-domain interactions and receptor multimerization suggest that ectodomain elements apart from the ligand-binding Ig1 domain may be important for Slit response and repulsive signaling *in vivo*. We have previously shown that Robo1 Ig1 is the only domain that is indispensable for Slit binding and midline repulsive signaling [21,22,43], but functional redundancy among other Robo1 ectodomain elements remains a possibility. Here, we identify the minimal ectodomain structure necessary for proper midline repulsive signaling by Robo1: we show that the Ig2-5 and Fn1-3 regions are dispensable for Slit binding and midline repulsion, and Ig1 alone is sufficient for a reduced level of midline repulsive signaling. If one other native domain is added back in addition to Ig1, normal repulsive activity is fully restored. The identity of the second domain appears unimportant, suggesting a permissive role for domains other than Ig1. Our results suggest that Robo receptor multimerization and ectodomain structural configurations observed *in vitro* may not be important for *in vivo* signaling, at least in the context of midline axon repulsion. We also report a novel role for the Robo1 Fn repeats in Robo2 trans-inhibition, providing further insight into the spatio-temporal regulation of Robo1 activity during midline crossing of axons in the embryonic central nervous system.

## Results

### Robo1’s Ig2-5 and Fn1-3 regions are dispensable for Slit binding and midline repulsion

We have previously reported that Ig1 is the only ectodomain element of Robo1 that is indispensable for Slit binding and midline repulsion [21,22,43]. We next sought to determine if the remaining Igs (Ig2-5) or Fns (Fn1-3) possess a redundant structural characteristic that can compensate for the loss of another to effectively bind Slit *in vitro* and signal midline repulsion *in vivo*. To answer this, we first tested whether deleting domains Ig2-5 or Fn1-3 together would impede Robo1’s ability to bind Slit in cultured *Drosophila* S2R+ cells. We transfected S2R+ cells with HA-tagged Robo1 variants (Robo1ΔIg2-5 or Robo1ΔFn1-3), then treated these cells with Slit-expressing media and stained with anti-HA and anti-Slit to recognize the Robo1 variants expressed within the cells and the Slit bound to the cell surface, respectively. Both Robo1ΔIg2-5 and Robo1ΔFn1-3 were able to bind Slit and were properly localized to the plasma membrane to the same degree as full-length Robo1 (Fig. 1B,C,D), demonstrating that the Ig2-5 and Fn1-3 regions of Robo1 are not required for Slit binding at the membrane of cultured *Drosophila* cells.

**Figure 1.**
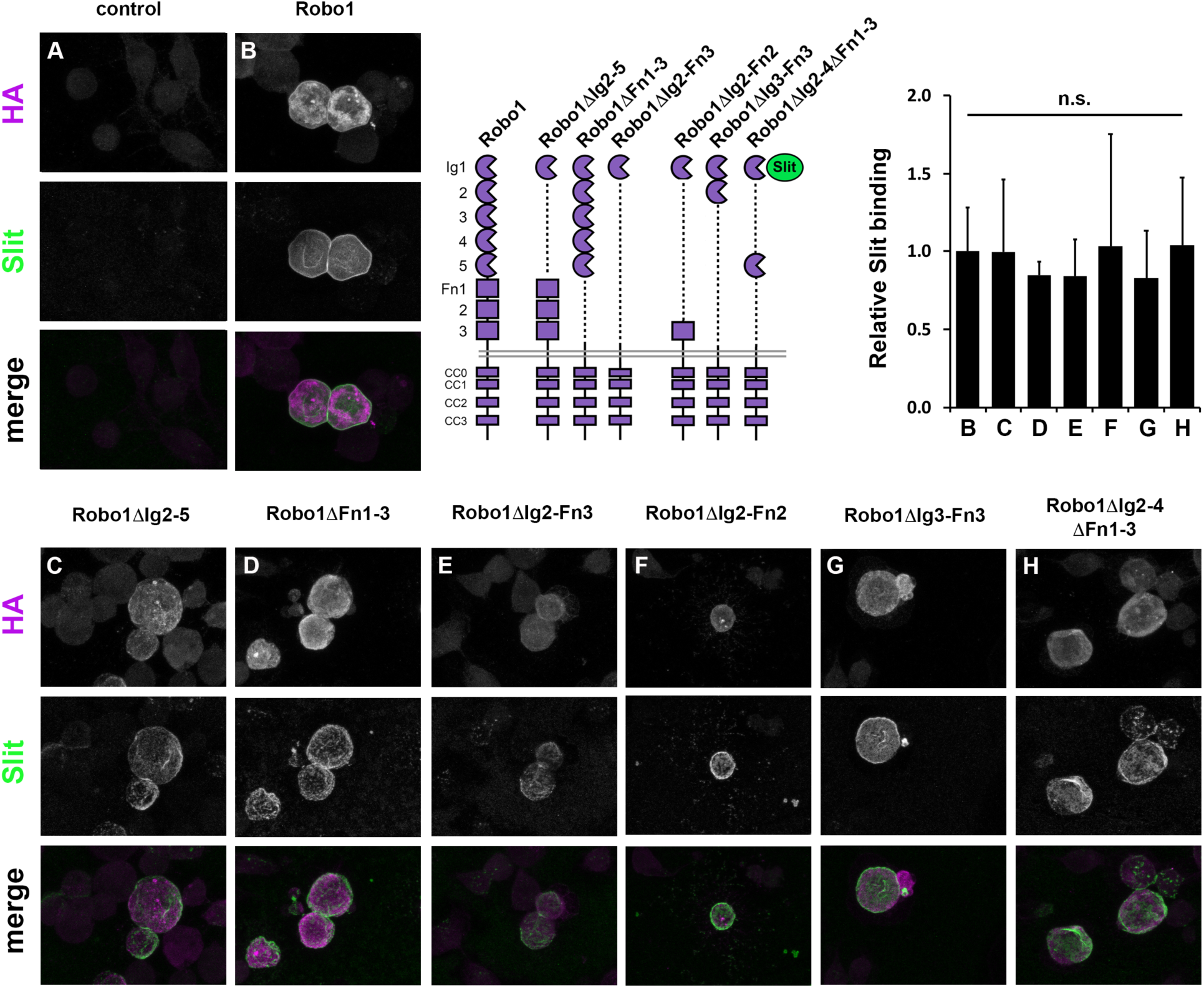
Robo1 variants containing an Ig1 domain can bind slit *in vitro*. Cultured *Drosophila* S2R+ cells expressing full-length Robo1 (B) or a version of Robo1 with elements of its ectodomain deleted (Robo1ΔIg2-5 C, Robo1ΔFn1-3 D, Robo1ΔIg2-Fn3 E, Robo1ΔIg2-Fn2 F, Robo1ΔIg3-Fn3 G, Robo1ΔIg2-4Fn1-3 H) were treated with media containing Slit. After Slit treatment, Robo1-expressing cells were fixed and stained with antibodies to detect HA-tagged Robo1 (magenta), and Slit (green). Slit binds to cells expressing full-length Robo1 and deletion variants C-H, but does not bind to untransfected cells (A). Receptor schematics show full-length Robo1 and Robo1 variant deletion constructs. Bar graph quantifies Slit binding levels in conditions B-H, normalized to the level observed with full-length Robo1 (B). Error bars show s.d. No statistical difference in Slit binding was detected for any Robo1 variant (C-H) compared to full-length Robo1 by Student’s two-tailed t-test (p>0.9 for all variants tested).

To test our deletion variants *in vivo* we used a genomic rescue construct in which HA-tagged variant *robo1* cDNAs are cloned into a plasmid containing regulatory sequence from the endogenous *robo1* gene (Fig. 2A) [21,22,44]. These plasmids also contain an attB site to allow FC31-directed site-specific integration into attP landing sites at the same cytological location (28E7) to ensure equivalent expression between transgenes. In wild-type embryos, endogenous Robo1 protein is detectable at high levels on longitudinal axons but present only at low levels on commissural axons (we refer to this restriction of expression pattern as “commissural clearance”) [21,38]. Transgenic HA-tagged Robo1 protein and Robo1ΔIg2-5 expressed from our rescue construct reproduces this pattern (Fig. 2B,C). In contrast, Robo1ΔFn1-3 is present on commissures in addition to longitudinals (Fig. 2D). This result is consistent with our previous study which indicated Robo1 Fn3 to be required for exclusion of Robo1 from commissural axons [43]. This data provides further evidence that Fn3 has a role in preventing Robo1 from reaching the growth cone surface in midline-crossing commissural axons, and/or in maintaining its clearance from commissures after midline crossing.

**Figure 2.**
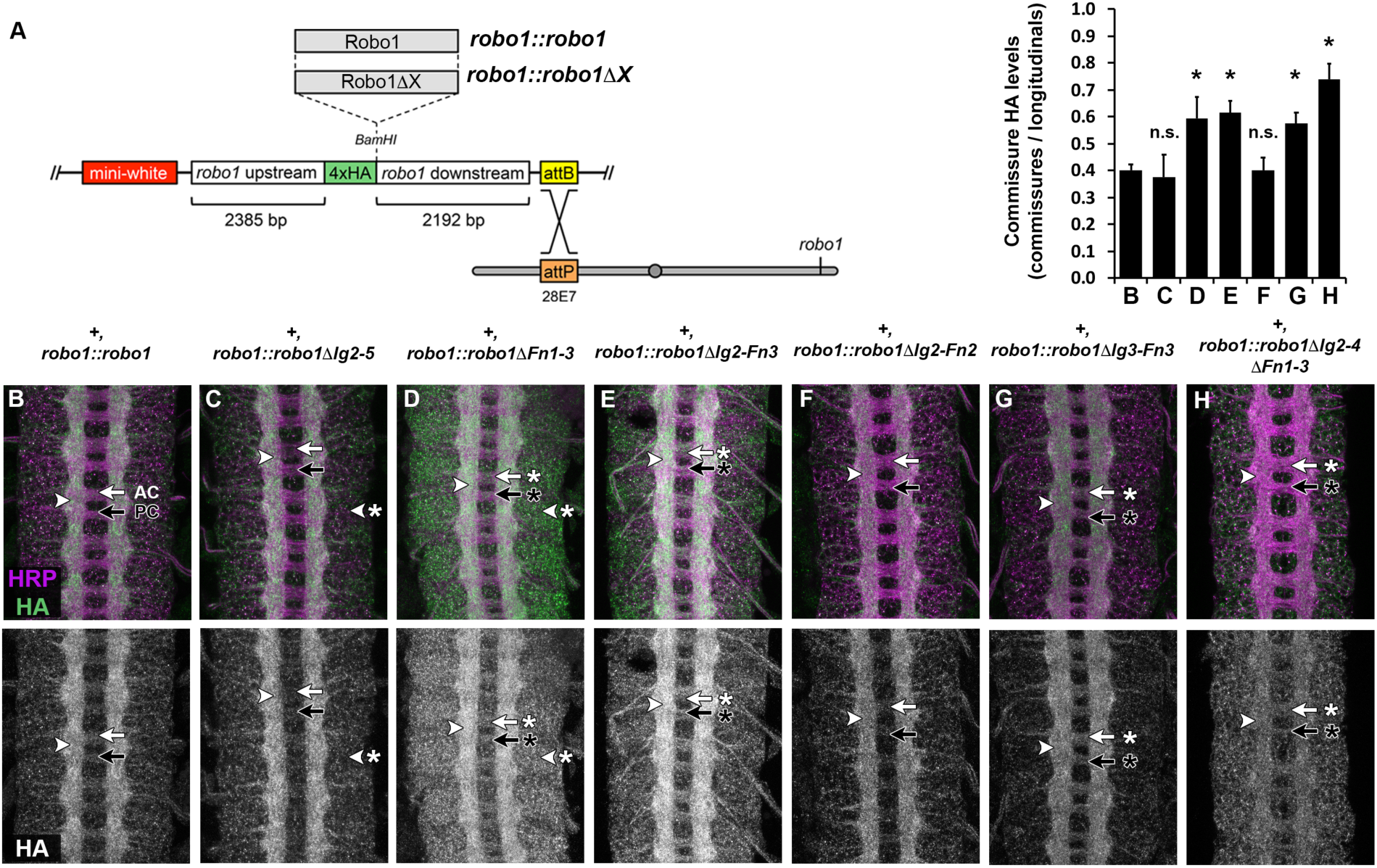
Robo1 Fn3 is required for commissural clearance. Expression of full-length Robo1 and Robo1 variants via the *robo1::robo1* genomic rescue transgene in a wild type background. (A) Schematic of robo1 rescue construct. (B-H) Stage 16 embryos stained with anti-HRP (magenta) and anti-HA (green) antibodies. Bottom images show HA channel alone from the same embryos. HA tagged full-length Robo1 (B), Robo1ΔIg2-5 (C), Robo1ΔFn1-3 (D), Robo1ΔIg2-Fn3 (E), Robo1ΔIg2-Fn2 (F), Robo1ΔIg3-Fn3 (G), or Robo1ΔIg2-4ΔFn1-3 (H) proteins expressed from the robo1 rescue transgene are localized to longitudinal axon pathways (arrowhead). However, variants lacking Fn3 [Robo1ΔFn1-3 (D), Robo1ΔIg2-Fn3 (E), Robo1ΔIg3-Fn3 (G) and Robo1ΔIg2-4ΔFn1-3 (H)] are present on commissural axon segments in the anterior commissure (AC, white arrow with asterisk) and posterior commissure (PC, black arrow with asterisk) to a higher degree than full-length Robo1 (B), Robo1ΔIg2-5 (C) or Robo1ΔIg2-Fn2 (F). Elevated expression in neuronal cell bodies is seen in Robo1ΔIg2-5 (D) and Robo1ΔFn1-3 (E) (arrowhead with asterisk). Bar graph quantifies the ratio of HA levels on commissural axons vs longitudinal axons for the genotypes shown in B-H (error bars show s.d.). Extent of commissural clearance for each Robo1 variant (C-H) was compared to full-length Robo1 (B) by a two-tailed Student’s t-test with a Bonferroni correction for multiple comparisons (*p<0.0001 compared to *robo1::robo1*). n=3 embryos per genotype.

To investigate the ability of these combinatorial deletion variants to carry out the receptor’s midline repulsive function, we introduced these transgenes into a *robo1* null mutant background. We then stained with anti-HA to recognize localization of the transgene. In this background, expression of either variant (Robo1ΔIg2-5 or Robo1ΔFn1-3) was able to restore wild-type axon scaffold morphology to the same degree as full-length Robo1 (Fig. 3A,B,C). We stained these embryos with the longitudinal axon pathway marker anti-FasII to investigate the ability of these variants to rescue midline repulsive activity in the absence of endogenous *robo1* by quantifying ectopic crossing of FasII-positive axons. In wildtype embryos, these axons remain on their own side of the body and do not cross the midline (Fig. 4A). However, midline repulsion is lost in *robo1*^*1*^ null mutant embryos and FasII-positive axons cross and recross the midline ectopically in every segment (Fig. 4B). Expression of full-length Robo1 (Fig 4C), Robo1ΔIg2-5 (Fig 4D), or Robo1ΔFn1-3 (Fig 4E) was able to completely restore midline repulsive function comparable to wildtype embryos, demonstrating that Robo1 retains full midline repulsive activity when four out of five of its Ig domains (Ig2-5) or all three of its Fn domains (Fn1-3) are deleted.

**Figure 3.**
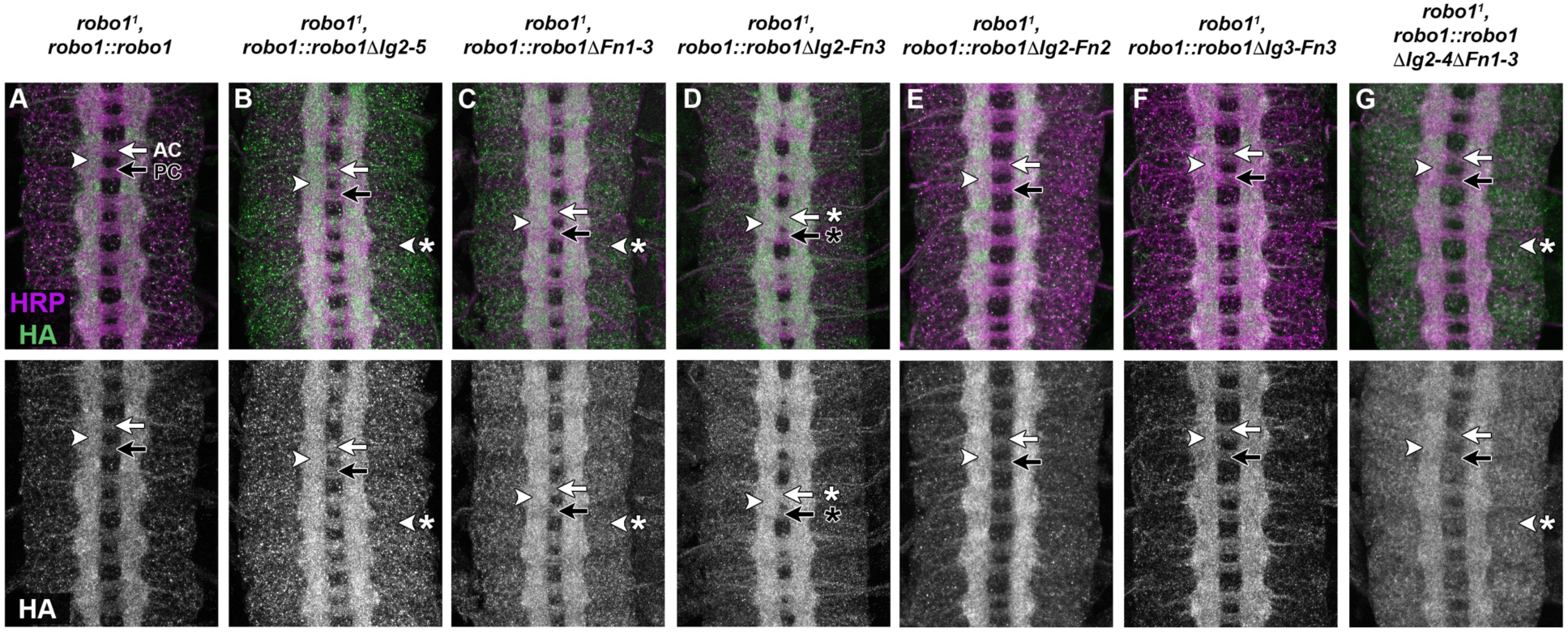
Expression of Robo1 variants in *robo1* mutant embryos. Stage 16 embryos stained with anti-HRP (magenta) and anti-HA (green) antibodies. Bottom images show HA channel alone from the same embryos. (A–G) Expression of full-length Robo1 via the *robo1* rescue transgene in a *robo1* null mutant (A) restores the wild-type structure of the axon scaffold. Each of the combinatorial Ig (ΔIg2-5) and Fn (ΔFn1-3) deletion variants restore axon scaffold morphology to a similar extent as full-length Robo1 (B-C). However, when the two variant deletions are combined (Robo1ΔIg2-Fn3) the transgene is not able to completely restore the scaffold (D, arrows with asterisk). Adding back one domain restores wild-type axon scaffold morphology (Robo1ΔIg2-Fn2 E, Robo1ΔIg3-Fn3 F, Robo1ΔIg2-4ΔFn1-3 G). In the absence of endogenous *robo1*, all of the variants are localized to the longitudinal pathways as in wild-type embryos (arrowheads). Robo1ΔIg2-5, Robo1ΔFn1-3, and Robo1ΔIg2-4ΔFn1-3 display elevated expression levels in neuronal cell bodies (B-C, G arrowhead with asterisk) compared to the other Robo1 variants.

**Figure 4.**
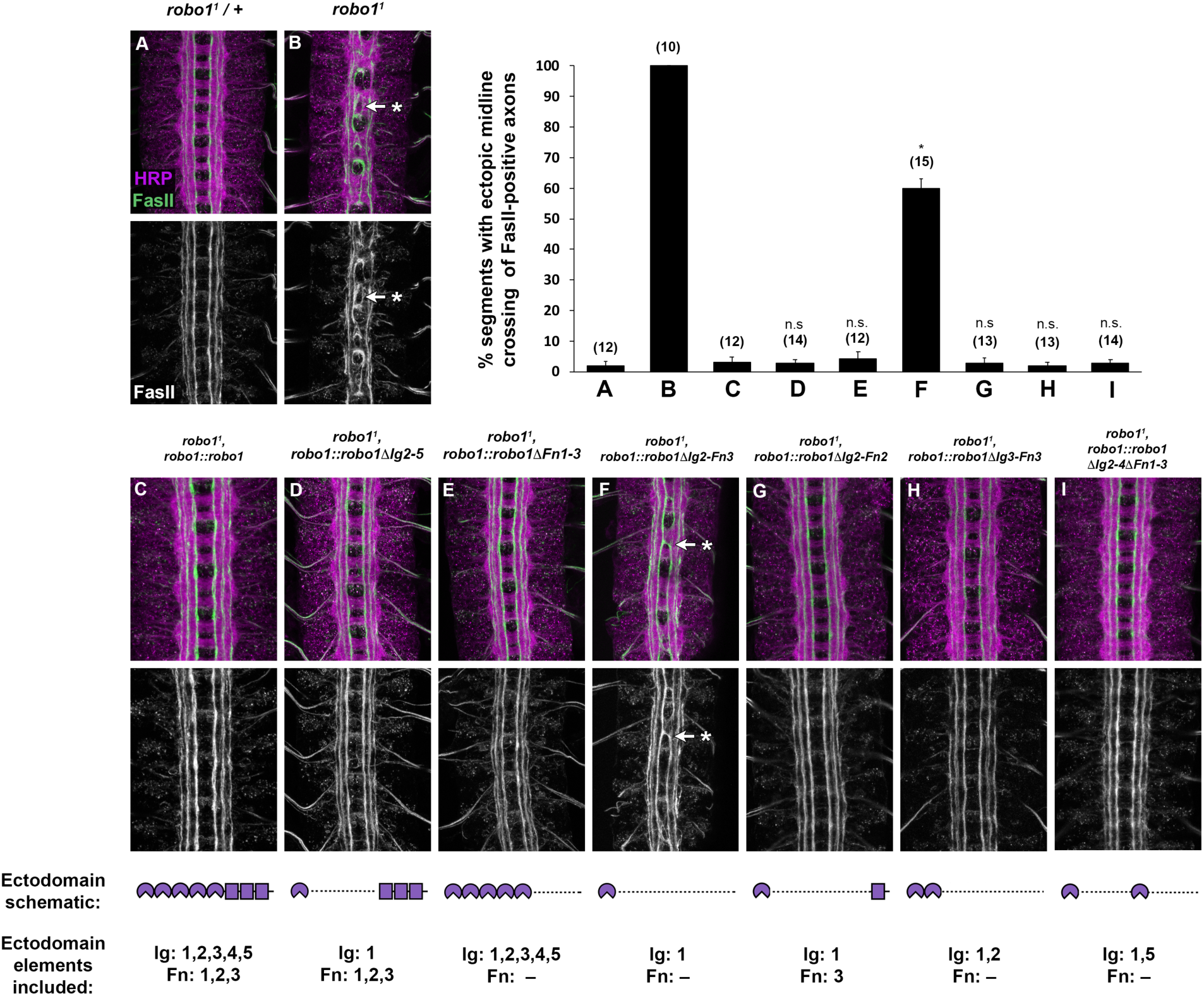
Ig1 alone can partially rescue Robo1’s midline repulsive activity. (A-I) Stage 16 embryos stained with anti-HRP (magenta) and anti-FasII (green) antibodies. Lower images show FasII channel alone for the same embryos. In wild-type embryos, FasII-positive axons do not cross the midline (A). However, in *robo1* null mutants, FasII-positive axons cross the midline inappropriately in every segment (B, arrow with asterisk). This phenotype is completely rescued by the robo1 rescue transgenes expressing full-length Robo1 (C) and Robo1 variant deletions Ig2-5 (D) and Fn1-3 (E) but is not completely rescued by the transgene expressing Robo1ΔIg2-Fn3 (F, arrow with asterisk). Although ectopic crossing is seen in sixty percent of abdominal segments in Robo1ΔIg2-Fn3 variants, the phenotype is less severe than *robo1* mutants in that the axons do not form characteristic roundabouts at the midline (compare F to B). However, the addition of one ectodomain (Fn3 G, Ig2 H, or Ig5 I) to Ig1 is enough to fully restore midline repulsion to wild-type levels. Bar graph at top right indicates instances of ectopic crossing in the genotypes shown in A-I with sample size of scored embryos (n) listed above each genotype. Error bars indicate s.e.m. The extent of rescue for each Ig deletion variant (D-I) was compared to *robo11, robo1::robo1* embryos (C) by a two-tailed Student’s t-test, with a Bonferroni correction for multiple comparisons (*p<0.01 compared to *robo11, robo1::robo1*).

### Ig1 alone is sufficient for partial rescue of midline repulsion in *robo1* null mutants

We next sought to determine whether a certain amount of the receptor’s ectodomain must be present for proper midline repulsive signaling, or whether the Slit-binding Ig1 domain alone is sufficient. To that end, we created a Robo1 variant in which all ectodomain elements except Ig1 are deleted (Robo1ΔIg2-Fn3) and tested its ability to bind Slit *in vitro* and signal midline repulsion *in vivo*. Cultured *Drosophila* S2R+ cells that express Robo1ΔIg2-Fn3 demonstrate co-localized staining at the plasma membrane to the same degree as full-length Robo1 or the other variants tested in this study (Fig. 1E). This indicates that Slit can still bind to the Robo1ΔIg2-Fn3 protein, which contains only Ig1 in its ectodomain.

We introduced the HA-tagged Robo1ΔIg2-Fn3 to embryos via our previously described genomic rescue construct and monitored expression levels and localization of the Robo1 variant in embryonic neurons. Consistent with our other Robo1 variants lacking the third Fn repeat, Robo1ΔIg2-Fn3 was localized primarily to longitudinal axons, but was also detectable on commissures (Fig. 2E).

When introduced into a *robo1* null mutant background, the Robo1ΔIg2-Fn3 transgene was not able to completely rescue the axon scaffold’s morphology, and *robo1*^*1*^, *robo1::robo1ΔIg2-Fn3* embryos exhibit defects that resemble a partial loss of *robo1* function, including thickening and partial fusion of commissures (Fig. 3D). We also observed increased HA staining on commissural axons in these embryos, consistent with ectopic midline crossing of longitudinal axons. However, the defects in these embryos do not appear as severe as in *robo1*^*1*^, *robo1::*robo1ΔIg1 embryos [21], suggesting the Robo1ΔIg2-Fn3 variant is phenotypically intermediate between the wildtype and *robo1* null mutants. When stained with anti-FasII, *robo1*^*1*^, *robo1::robo1ΔIg2-Fn3* embryos show FasII-positive axons crossing in 60% of abdominal segments, compared to 100% in *robo1*^*1*^ null mutants (see Fig 4 bar graph for quantification). Our quantitative and qualitative observation of the ectopic crossing in these embryos alongside the expression pattern of our HA-tagged Robo1ΔIg2-Fn3 variant in a *robo1* null mutant background indicate this variant containing just Ig1 can partially rescue midline repulsive function.

### Robo1 variants containing Ig1 plus one additional native domain exhibit midline repulsive activity equivalent to full-length Robo1

While Ig2-5 and Fn1-3 are each dispensable for midline repulsive function, deleting both of these regions so that only Ig1 remains reduces Robo1’s ability to signal midline repulsion. To determine the minimal number of ectodomain elements required for proper signaling, we constructed three additional variants where one native ectodomain element was added back in addition to Ig1: Robo1ΔIg3-Fn3 (ectodomain contains only Ig1+Ig2), Robo1ΔIg2-4ΔFn1-3 (ectodomain contains only Ig1+Ig5), and Robo1ΔIg2-Fn2 (ectodomain contains only Ig1+Fn3) (see Figs. 1 and 4 for Robo1 variant schematics). Each of these Robo1 variants displayed Slit binding levels equivalent to full-length Robo1 in our S2R+ Slit binding assay (Fig. 1F-H).

When expressed in embryonic neurons via our *robo1* rescue construct, we found that Robo1ΔIg2-Fn2 and Robo1ΔIg3-Fn3 both were expressed at similar levels to full-length Robo1 and localized to longitudinal axons (Fig. 2B,F,G). However, Robo1ΔIg2-4ΔFn1-3 was expressed at lower levels than the other variants or full-length Robo1 (Fig. 2H). This could potentially be due to this construct containing two artificial junctions, one between Ig1 and Ig5 and the other between Ig5 and the juxtamembrane region, as opposed to the one artificial junction present in all previously described constructs. We also note that variants lacking Fn3 (Robo1ΔIg3-Fn3 and Robo1ΔIg2-4ΔFn1-3) remain on commissures. We have now described five variants lacking Fn3 (Robo1ΔFn3 [43], Robo1ΔFn1-3, Robo1ΔIg2-Fn3, Robo1ΔIg3-Fn3, and Robo1ΔIg2-4ΔFn1-3) in which Robo1 protein is present at elevated levels on commissures, consistent with a requirement for Fn3 of Robo1 for the receptor’s commissural clearance.

We next introduced our Robo1 transgenes into a *robo1* null mutant background to assay if adding back an additional domain would restore their ability to rescue midline repulsive activity when endogenous *robo1* is absent. We found that expression of any of the variants containing Ig1 plus one other ectodomain element were able to fully restore wild-type axon scaffold morphology (Fig. 3E,F,G). Quantification of FasII ectopic crossing in these variants confirmed a complete rescue of midline repulsion (Fig. 4G,H,I). This indicates that at least six of the eight ectodomain elements can be deleted without affecting Robo1’s ability to signal midline repulsion, and signaling efficiency does not seem to rely upon the type of domain present apart from Ig1, as Ig2, Ig5, and Fn3 show similar rescue when combined with Ig1 in our minimal ectodomain variants.

### Robo1 variants lacking Fn domains are insensitive to Comm downregulation

In *Drosophila*, Commissureless (Comm) serves as a negative regulator to the Slit-Robo1 pathway by preventing newly-synthesized Robo1 protein from reaching the surface of axonal growth cones. This allows axons to cross the midline and innervate a target on the opposite side of the body [37-39]. To determine whether Robo1’s ectodomain elements are collectively dispensable for Comm-dependent regulation, we used the GAL4/UAS system to force high levels of ectopic Comm expression in embryos carrying each of the previously described Robo1 deletion variants (*robo1::robo1ΔIg2-5, robo1::robo1ΔFn1-3*, and *robo1::robo1ΔIg2-Fn3*) and observed the expression and localization of the Robo1 variants within the embryonic nerve cord by using anti-HA.

Forcing pan-neural Comm expression in embryos encourages a *slit*-like axon scaffold collapse and the strong downregulation of HA-tagged Robo1 variants on axons [21,22,38,43]. Consistent with previous results, Comm-dependent downregulation of Robo1 expression does not depend on its Ig domains, as we observed a strong reduction in neuronal HA staining in embryos carrying Robo1ΔIg2-5 along with *elav-GAL4* and *UAS-Comm*, as well as thickened commissures consistent with an increase in midline crossing due to down-regulation of endogenous Robo1 (Fig. 5F). However, when the Fn3 repeat is absent in either Robo1ΔFn1-3 or Robo1ΔIg2-Fn3 transgenes, the variant protein remains on axons of UAS-Comm embryos (compare Fig. 5 C and G; D and H). The strong midline collapse phenotype caused by Comm misexpression in embryos expressing Robo1ΔFn3 [43], Robo1ΔFn1-3 and Robo1ΔIg2-Fn3 suggests that Comm retains the ability to antagonize these proteins through a non-sorting mechanism, as has previously been described for sorting-deficient forms of Robo1 [39].

**Figure 5.**
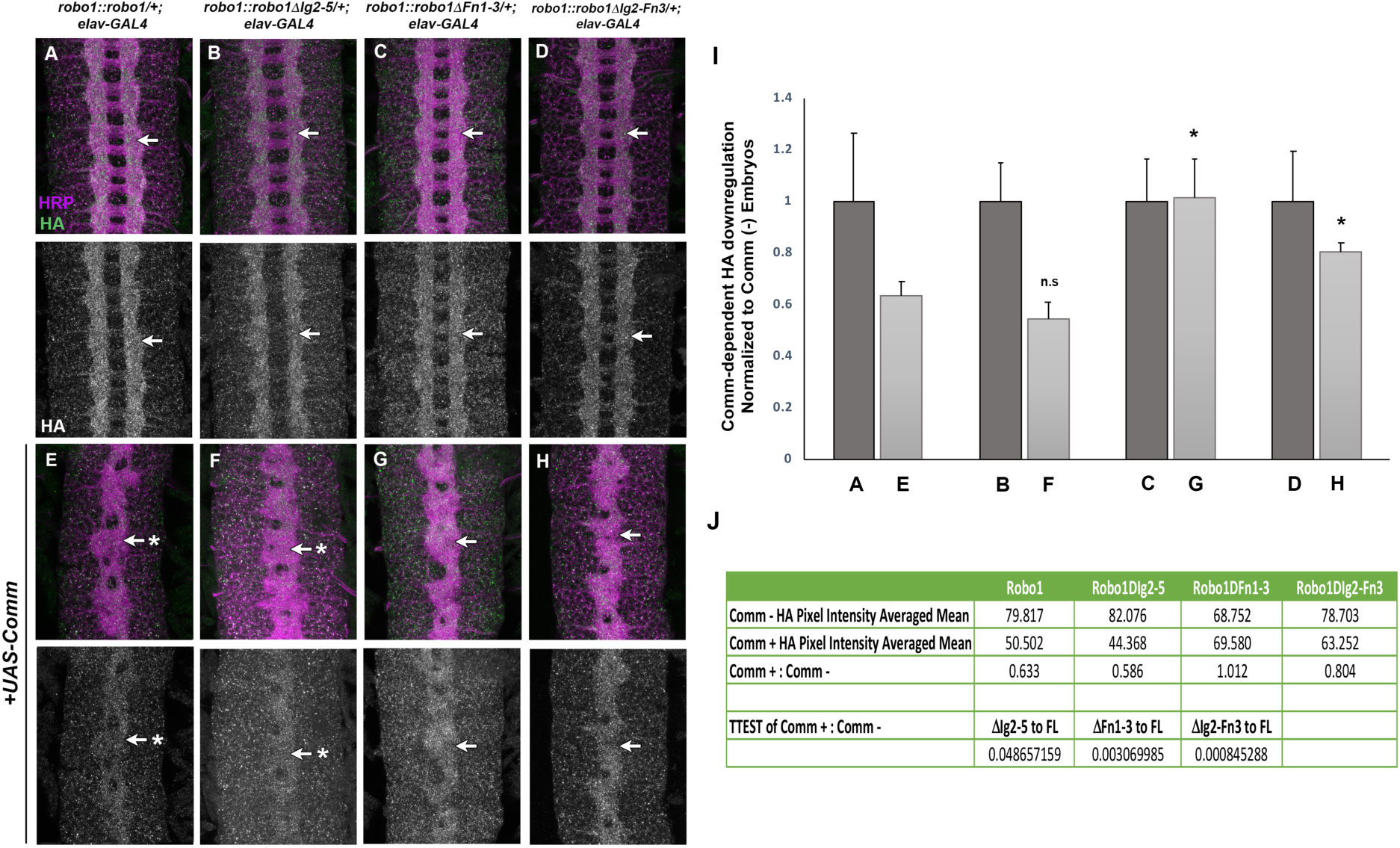
Robo1’s Fn3 domain is required for Comm-dependent downregulation. Stage 16 embryos stained with anti-HA (green) and anti-HRP (magenta). Lower images show HA channel alone of the same embryos. (A-D) Embryos with one copy of the transgene as well as elav-GAL4 display normal Robo1 protein expression among the HA-tagged variants (arrows). (E-H) Homozygous transgenic embryos carrying elav-GAL4 and UAS-Comm show strongly downregulated HA expression among the slit-like collapsed axon scaffold (arrows with asterisks), with the exceptions of Robo1ΔFn1-3 and Robo1ΔIg2-Fn3 (G and H) which are not downregulated on axons when Comm is misexpressed (arrow). Pairs of sibling embryos shown (A and E; B and F; C and G; D and H) were stained in the same tube and imaged under the same confocal settings to ensure accurate comparison of HA levels between embryos. Bar graph (I) and table (J) show quantification of pixel intensity of full-length and combinatorial Robo1 variants with either elav-GAL4 alone (A-D) or elav-GAL4 and Comm (E-H). Error bars indicate s.d. For each of these variants, the degree of HA downregulation was compared to that of full-length Robo1 by a two-tailed Student’s t-test, with a Bonferroni correction for multiple comparisons (*p<0.01). n=5 for each UAS-Comm and elav-GAL4 genotype.

### Robo1 Fn domains are necessary for negative regulation by Robo2

A second Robo family member in *Drosophila*, Robo2, plays a dual role in both promoting and inhibiting midline crossing in the embryonic CNS. Robo2 is co-expressed with Robo1 in ipsilateral pioneer neurons, where they cooperate to prevent midline crossing in response to Slit [45,46]. Robo2 is also expressed in midline cells during early stages of axon guidance, where it interacts with Robo1 in trans to inhibit Robo1’s response to Slit and promote midline crossing [29]. This negative regulatory role of Robo2 depends on its Ig1 and Ig2 domains, but it is unknown which region(s) of Robo1 contribute to negative regulation by Robo2. Our collection of Robo1 deletion variants provided us an opportunity to map the critical sequences in Robo1 that confer sensitivity to Robo2 inhibition. To this end, we used *elav-GAL4* and *UAS-Robo2* transgenes to misexpress Robo2 at high levels in all neurons (and transiently in midline glia) in embryos expressing our Robo1 deletion variants in place of endogenous *robo1* (Fig. 6).

**Figure 6.**
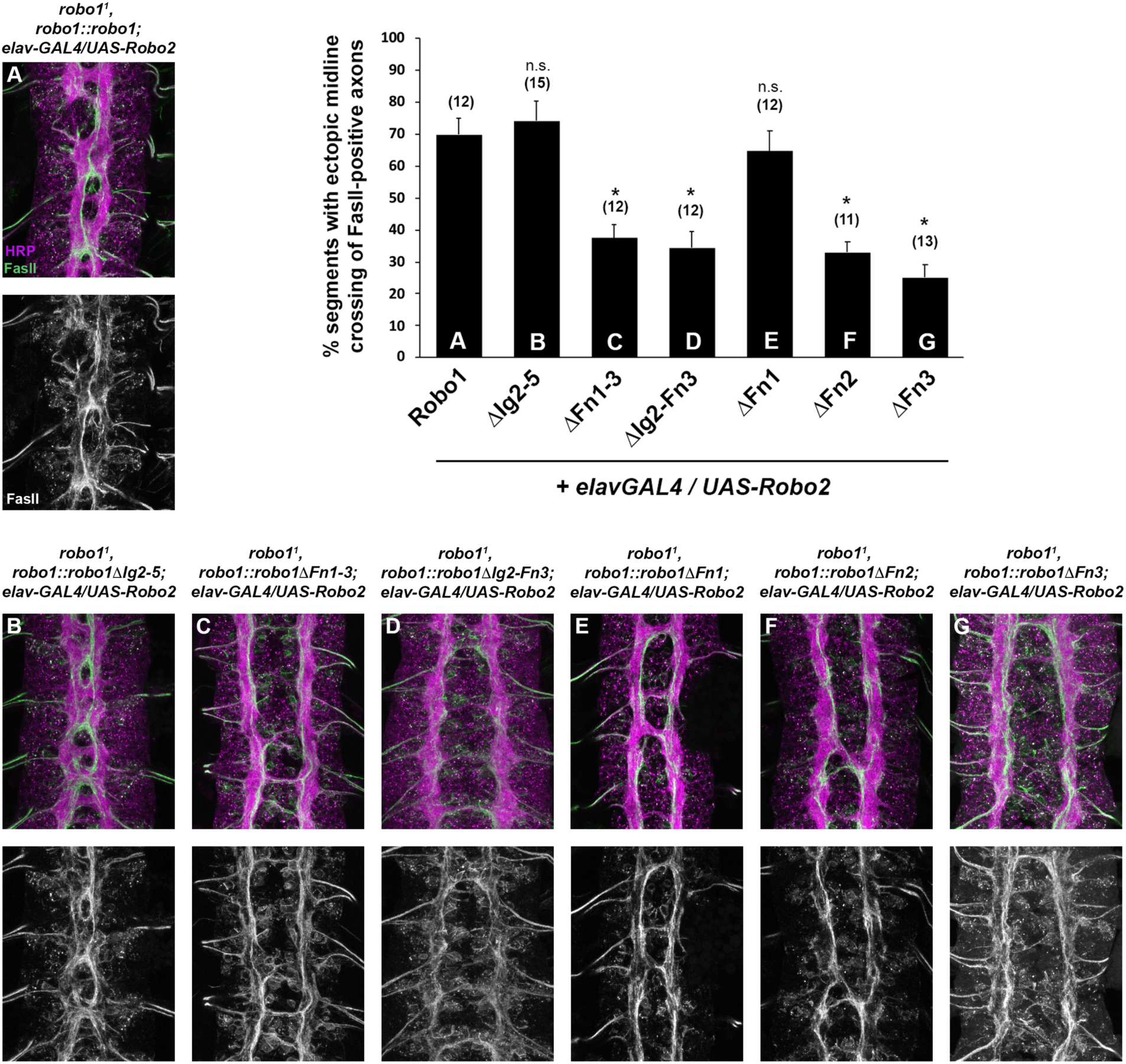
The Fn2 and Fn3 repeats of Robo1 are required for sensitivity to Robo2. Expression of Robo1 and Robo1 variants in a Robo2 misexpression background. Stage 16 embryos stained with anti-HRP (magenta) and anti-FasII (green) antibodies. Misexpression of Robo2 (via *elav-GAL4* and *UAS-Robo2*) induces strong ectopic midline crossing of FasII-positive axons in embryos that express full length Robo1 (A), Robo1ΔIg2-5 (B), or Robo1ΔFn1 (E). However, Robo2 misexpression in embryos expressing Robo1ΔFn1-3 (C), Robo1ΔIg2-Fn3 (D), Robo1ΔFn2 (F), or Robo1ΔFn3 (G) instead display a commissureless phenotype with significantly decreased levels of ectopic midline crossing. Qualitatively, the phenotypic differences observed suggest that Robo1 variants lacking any of the Fn repeats are insensitive to transient downregulation by Robo2. Bar graph at top left indicates instances of ectopic crossing in the genotypes shown in (A-G) with sample size of scored embryos (n) listed above each genotype. Error bars indicate s.e.m. Ectopic midline crossing defects caused by Robo2 misexpression in each Robo1 variant background (B-G) were compared to those caused by Robo2 misexpression in a full-length Robo1 background (A) by a two-tailed Student’s t-test, with a Bonferroni correction for multiple comparisons (*p<0.0001).

Pan-neural misexpression of Robo2 induces a strong ectopic midline crossing phenotype in embryos expressing endogenous Robo1, as Robo2’s inhibition of Slit-Robo1 repulsion has a stronger effect than its own ability to induce ectopic repulsion [29,33,45]. Robo2 variants that are unable to interact with Robo1 (for example Robo2ΔIg2) instead induce ectopic repulsion, as only Robo2’s midline repulsive activity remains intact in this context [29].

We found that misexpressing Robo2 in *robo1* mutants carrying our full-length *robo1* rescue construct *(robo1*^*1*^,*robo1::robo1; elav-GAL4/UAS-Robo2)* produced a strong ectopic crossing phenotype similar to the effect of misexpressing Robo2 in a wild type background (Fig. 6A) [29]. We observed an identical effect when Robo2 was misexpressed in *robo1*^*1*^,*robo1::robo1ΔIg2-5* embryos, indicating that Robo1ΔIg2-5 remains sensitive to inhibition by Robo2 (Fig. 6B). In contrast, both the Robo1ΔFn1-3 and Robo1ΔIg2-Fn3 variants were insensitive to Robo2 inhibition, as Robo2’s ability to induce ectopic crossing in these backgrounds was significantly reduced (Fig. 6C,D). Instead, Robo2 misexpression in these backgrounds produced a strongly commissureless phenotype, reflecting a shift away from Robo2-dependent midline crossing (via inhibition of Robo1) and favoring Robo2-dependent midline repulsion (which does not involve Robo1 inhibition). These results suggest that one or more of the Fn repeats of Robo1 are critical to the protein’s ability to be inhibited by Robo2.

To determine which of its three Fn repeats are responsible for Robo1’s sensitivity to Robo2, we misexpressed Robo2 in embryos expressing each of our previously described Robo1 individual Fn deletion variants (Robo1ΔFn1, Robo1ΔFn2, Robo1ΔFn3) in place of endogenous *robo1* [43]. We found that Robo1ΔFn1 retained Robo2 sensitivity, similar to full-length Robo1 and Robo1ΔIg2-5 (Fig. 6E), while Robo1ΔFn2 and Robo1ΔFn3 were insensitive to Robo2 inhibition, similar to Robo1ΔFn1-3 and Robo1ΔIg2-Fn3 (Fig. 6F,G). We therefore conclude that the Robo1 Fn2 and Fn3 repeats play a unique role in the interactions required for Robo2-dependent inhibition of Robo1, while Robo1 Fn1 is dispensable for this regulation.

## Discussion

In this paper, we have examined the structural requirements for midline repulsive signaling by the *Drosophila* Robo1 axon guidance receptor, by identifying the minimal combination of structural elements needed in the receptor’s ectodomain for Slit binding and midline repulsive signaling. Using a series of Robo1 variants in which various combinations of ectodomain elements are deleted, we show that the Slit-binding Ig1 domain alone is sufficient for Slit binding and partial midline repulsive activity, while adding back one additional native domain in combination with Ig1 restores midline repulsive activity to levels that are indistinguishable from full-length Robo1. Notably, the Ig2, Ig5, or Fn3 domains of Robo1 can each restore wild type function in combination with Ig1, suggesting that the identity of the second ectodomain element is not critical for repulsive signaling, and that domains other than Ig1 may play a permissive role in Slit-dependent receptor activation. Our results also reveal a novel requirement for two Robo1 Fn domains (Fn2 and Fn3) for its regulation by Robo2.

### Ig1 alone is not sufficient for full midline repulsive signaling by Robo1

We have previously reported that each of *Drosophila* Robo1’s ectodomain elements except Ig1 are individually dispensable for midline repulsion *in vivo* [21,22,43]. Here we find that while Ig1 alone is not sufficient for full repulsive activity of Robo1, a Robo1 variant containing only Ig1 in its ectodomain (Robo1ΔIg2-Fn3) displayed reduced but significant levels of midline repulsive activity. Our Robo1ΔIg2-Fn2, Robo1ΔIg3-Fn3 and Robo1ΔIg2-4ΔFn1-3 variants all reintroduced one native domain of similar length (103AA, 100AA and 94AA, respectively) to our Robo1ΔIg2-Fn3 variant and were all able to restore midline repulsion to wild-type levels. As the levels of rescue do not depend on the identity of the second ectodomain element, or even its domain type (Ig or Fn), one possibility is that Ig1 alone is sufficient for Slit binding and receptor activation *in vivo* as long as it is maintained at a minimum distance from the membrane. In this case, the partial rescue seen with Robo1ΔIg2-Fn3 might reflect a decrease in Robo1’s ability to bind Slit due to steric hindrance caused by Ig1’s proximity to the plasma membrane, while adding back one additional domain might be sufficient to increase the Ig1-membrane distance sufficiently to allow full access to Ig1 by Slit.

### Robo1 multimerization and mechanism of Robo1 activation

Previous interaction studies have shown that Slit is able to form a dimer when binding to Robo1 [18,47] and recent structural analysis of human Robo1 suggests that the receptor exists as an inactive dimer on the cell surface, via contacts primarily between Ig3 as well as Ig1 and Ig4, with the Ig1-4 region adopting an extended conformation state {Aleksandrova:2018dl}. This study further suggests that the oligomeric state of Robo1 may not change upon Slit2 binding, indicating that monomer-to-oligomer transition is not part of the mechanism of Robo1 activation. Importantly, this study did not address whether multimerization of Robo1 was necessary for Slit2 binding or receptor activation. Assuming *Drosophila* Robo1 and human Robo1 are activated via the same mechanism, our *in vivo* evidence suggests that Robo1 multimerization is not a pre-requisite for Slit binding or receptor activation, as homophilic interactions would presumably be compromised in our Robo1 variants lacking Ig3-4. Our Robo1ΔIg2-Fn3 Slit binding results indicate that this protein can still bind Slit (Fig. 1E). Our use of anti-SlitC (which recognizes the C-terminal of Slit) in this experiment detects binding between Robo1 and full-length Slit, but not Slit-N, and does not reveal the oligomerization state of the bound Slit. If close association of two or more Robo1 molecules is necessary for receptor activation, and Robo1 variants lacking Ig3-4 are unable to multimerize on their own, perhaps a pre-formed Slit dimer may be able to bridge two Robo1 monomers and facilitate their interactions independently of Ig3-4.

Upon Slit binding, Robo1 is subject to regulated proteolytic cleavage by the ADAM family metalloprotease Kuzbanian (Kuz) and Clathrin-dependent endocytosis, and both of these steps appear to be required for Slit-dependent midline repulsive signaling [23,24]. Although the precise location of Robo1’s Kuz cleavage site remains unknown, it is likely to be present in our Fn deletion variants, which retain the 46 aa juxtamembrane region between Fn3 and the transmembrane domain. A chimeric receptor in which all three Fn domains and the juxtamembrane region of Robo1 were replaced by equivalent sequences from Frazzled was resistant to Kuz cleavage and unable to rescue midline crossing defects in *robo1* mutants [23]; considering that deleting Fn1-3 does not detectably impair midline repulsion in our assays, sequences within the three Fn domains are unlikely to be required for Kuz cleavage if this is an obligate step in Robo1 activation.

### Fn domain-dependent regulation by Robo2 and Commissureless

Our results indicate that Robo1 Fn domains are required for negative regulation by Comm (Fn3 only) and Robo2 (Fn2 and Fn3). As Robo2 regulates Robo1 via trans binding interactions [29], the observation that deleting either Robo1 Fn2 or Fn3 domains reduces its sensitivity to Robo2 inhibition may indicate multiple points of contact between Robo1 and Robo2, or perhaps that these two Fn domains are required for a specific conformation of Robo1 that facilitates Robo2 binding. The requirement of multiple domains for protein-protein interactions has been previously described not only in the Robo2 domains (Ig1, Ig2) required for inhibitory Robo1-Robo2 heterodimers but also in the *in vitro* formation of receptor homodimers as seen in Dscam (Ig2, Ig3, Ig7) and Robo1 (Ig1, Ig3, Ig4) [29,48]. It will be interesting to characterize the exact binding points between Robo2 Ig1-2 and Robo1 Fn2-3 as well as to determine how exactly Robo2 binding inhibits Robo1 activation.

Fn3 is required for the receptor’s downregulation by both Comm and Robo2. If variants lacking Fn3 are insensitive to downregulation, why would they not exhibit enhanced midline repulsion? We note that phenotypic defects caused by loss of Robo2-dependent inhibition of Robo1 are only evident when attractive Net-Fra signaling is also compromised, for example in *robo2,fra* compound mutants [29,44]. Comm can also inhibit Slit-Robo repulsion independently of Robo sorting [39], and this sorting-independent regulation remains intact in all of our variants (including Robo1ΔIg2-Fn3; Fig. 3D). Thus we would not necessarily expect Robo1 variants that are insensitive to Comm sorting and Robo2 inhibition to display hyperactive midline repulsive activity in embryos with wild-type levels of Comm, Netrin, and Fra.

### Roles and requirements for Ig and Fn domains in other axon guidance receptors

The 5 Ig + 3 Fn ectodomain structure characteristic of Robo family receptors is broadly conserved, and most Robo receptors in most species share this 5 + 3 arrangement. The only known Robo family members to deviate from this characteristic structure are present in the silkworm, *Bombyx mori* (BmRobo1a and BmRobo1b), and in vertebrates (Robo4/Magic Roundabout) – where BmRobo1a/b are missing Ig5 and Fn1, and Robo4 is missing Ig3-5 and Fn1 [13,49]. In the case of *Bombyx* Robo1 paralogs, functional evidence suggests that these receptors play a canonical role in Slit-dependent midline repulsion during silkworm embryonic development, but Slit binding has not been directly examined [49]. Robo4 has roles in angiogenesis and neuronal migration and does not appear to be involved in midline repulsion [13,50,51]. While some reports indicate that Robo4 can bind Slit ligands [50], others suggest that Robo4 does not interact with Slit [52] but rather can act as a binding partner for the UNC5B receptor via Robo4’s Ig1-Ig2 region [15].

While we have shown here that the majority of the *Drosophila* Robo1 ectodomain is dispensable for midline repulsive signaling in the embryonic CNS, additional known or unknown roles of Robo1 may require structural features apart from Ig1. In other Robo family members, domains other than Ig1 have been implicated in diverse axon guidance contexts. In *Drosophila* Robo2, the Ig2 domain is required for trans-inhibition of Robo1 in commissural axons [29], and the Ig3 domain contributes to Robo2’s role in longitudinal pathway formation, possibly by promoting multimerization of Robo2 [33]. The mammalian Robo3/Rig-1 receptor has lost its ability to bind Slit due to amino acid substitutions in its Ig1 domain [14], but has acquired a novel ligand, NELL2, which binds to one or more of Robo3’s Fn domains [12]. Expression of NELL2 in the motor column of the spinal cord repels Robo3-expressing commissural axons away from lateral regions and towards the ventral midline, representing a unique role of Robo3 that does not appear to be shared among other Robo family members [12].

Outside of the Roundabout family, a number of other axon guidance receptor families have ectodomains that consist of Ig domains and Fn repeats, such as Frazzled/DCC (4 Ig + 6 Fn), Dscam (10 Ig + 6 Fn), and dLar (3 Ig + 8 Fn), among others. In some cases, biochemical studies have identified individual domains or combinations of domains that mediate interactions with ligands, receptor multimerization or homophilic trans binding, or interactions with other receptors. For example, homophilic interactions between *Drosophila* Dscam molecules depend on intermolecular contacts between Ig2, Ig3, and Ig7 [48,53], while Dscam can also interact with Slit-N via at least two redundant sites in the Ig1-5 region [47,54]. Vertebrate Dscam can also interact with Netrin ligands via Ig7-9 [55]. Interactions between the Frazzled/DCC receptor and its Netrin ligands are mediated by the Fn4 and Fn5 domains of DCC [56-58], while binding of another ligand, Draxin, is mediated by Ig1-Ig4 [59]. A heteromultimeric complex forms between vertebrate DCC and Robo3 receptors via interactions mediated by the cytoplasmic P3 domain of DCC and CC2+CC3 motifs of Robo3 [14]. However, few studies have systematically examined the functional requirements for individual domain elements via engineered domain deletions or sought to determine the minimal complement of structural elements necessary for *in vivo* signaling activity. Along with our previously reported structure/function studies of Robo1, the current study demonstrates that such studies are feasible and can provide insight into the signaling mechanisms and regulation of axon guidance receptors.

### Materials and methods

#### Molecular biology

*Robo1 variant deletions:* Robo1 domain deletions were generated via site-directed mutagenesis using Phusion Flash PCR MasterMix (Thermo Scientific), and completely sequenced to ensure no other mutations were introduced. Robo1 deletion variants include the following amino acid residues, relative to Genbank reference sequence AAF46887: Robo1ΔFn1 (Q52-P534/I646-T1395); Robo1ΔFn2 (Q52-T645/Y763-T1395); Robo1ΔFn3 (Q52-T762/H866-T1395); Robo1ΔIg2-5 (P56-V152/G535-T1395); Robo1ΔFn1-3 (P56-P534/H866-T1395); Robo1ΔIg2-Fn3 (P56-V152/H866-T1395); Robo1ΔIg2-Fn2 (P56-V152/Y763-T1395); Robo1ΔIg3-Fn3 (P56-Q252/H866-T1395); Robo1ΔIg2-4ΔFn1-3 (P56-V152/E441-P534/H866-T1395). Fn domain annotation shown in [43].

*pUAST cloning*: *robo1* coding sequences were cloned as BglII fragments into p10UASTattB for S2R+ cell transfection. All *robo1* p10UASTattB constructs include identical heterologous 5′ UTR and signal sequences (derived from the *Drosophila wingless* gene) and an N-terminal 3xHA tag.

*robo1 rescue construct cloning:* Construction of the *robo1* genomic rescue construct was described previously [21,22,43]. Full-length and variant Robo1 coding sequences were cloned as BglII fragments into the BamHI-digested backbone. Robo1 proteins produced from this construct include the endogenous Robo1 signal peptide, and the 4xHA tag is inserted directly upstream of the first Ig domain.

### Genetics

The *robo1*^*1*^ (also known as *robo*^*GA285*^) *Drosophila* mutant allele was used. The following *Drosophila* transgenes were used: *P{GAL4-elav*.*L}3 (elavGAL4), P{UAS-CommHA}*, P{10UAS-HARobo2}86Fb [29], *P{robo1::HArobo1}*, [21], *P{robo1::HArobo1ΔFn1}, P{robo1::HArobo1ΔFn2}, P{robo1::HArobo1ΔFn3}* [43], *P{robo1::HArobo1ΔIg2-5}, P{robo1::HArobo1ΔFn1-3}, P{robo1::HArobo1ΔIg2-Fn3}, P{robo1::HArobo1ΔIg2-Fn2}, P{robo1::HArobo1ΔIg3-Fn3}*, and *P{robo1::HArobo1ΔIg2-4ΔFn1-3}*. Transgenic flies were generated by BestGene Inc (Chino Hills, CA) using FC31-directed site-specific integration into attP landing sites at cytological position 28E7 (for *robo1* genomic rescue constructs). *robo1* rescue transgenes were introduced onto a *robo1*^*1*^ chromosome via meiotic recombination, and the presence of the *robo1*^*1*^ mutation was confirmed in all recombinant lines by DNA sequencing. All crosses were carried out at 25°C.

### Slit binding assay

*Drosophila* S2R+ cells were cultured at 25°C in Schneider’s media plus 10% fetal calf serum. To assay Slit binding, cells were plated on poly-L-lysine coated coverslips in six-well plates (Robo-expressing cells) or 75 cm^2^ cell culture flasks (Slit-expressing cells) at a density of 1-2×10^6^ cells/ml and transfected with pRmHA3-GAL4 [60] and HA-tagged p10UAST-Robo or untagged pUAST-Slit plasmids using Effectene transfection reagent (Qiagen). GAL4 expression was induced with 0.5 mM CuSO_4_ for 24 hours, then Slit-conditioned media was harvested by adding heparin (2.5 ug/ml) to Slit-transfected cells and incubating at room temperature for 20 minutes with gentle agitation. Robo-transfected cells were incubated with Slit-conditioned media at room temperature for 20 minutes, then washed with PBS and fixed for 20 minutes at 4°C in 4% formaldehyde. Cells were permeabilized with PBS+0.1% Triton X-100, then stained with antibodies diluted in PBS+2mg/ml BSA. Antibodies used were: mouse anti-SlitC (Developmental Studies Hybridoma Bank [DSHB] #c555.6D, 1:50), rabbit anti-HA (Covance #PRB-101C-500, 1:2000), Cy3-conjugated goat anti-mouse (Jackson #115-165-003, 1:500), and Alexa 488-conjugated goat anti-rabbit (Jackson #111-545-003, 1:500). After antibody staining, coverslips with cells attached were mounted in Aqua-Poly/Mount (Polysciences, Inc.). Confocal stacks were collected using a Leica SP5 confocal microscope and processed by Fiji/ImageJ [61] and Adobe Photoshop software. For quantification of Slit binding (Fig. 1), confocal max projections were opened in Fiji/ImageJ and individual cells were outlined using the “Freehand selections” tool. Pixel intensities for the anti-Slit and anti-HA (Robo1) channels were measured for 3-8 cells for each Robo1 variant, and the average ratio of Slit:HA for each construct (normalized to full-length Robo1) is reported as “relative Slit binding” in Fig. 1.

### Immunofluorescence and imaging

*Drosophila* embryo collection, fixation and antibody staining were carried out as previously described [62]. The following antibodies were used: FITC-conjugated goat anti-HRP (Jackson Immunoresearch #123-095-021, 1:100), Alexa Fluor 488-conjugated goat Anti-HRP (Jackson Immunoresearch #123-545-021, 1:500), mouse anti-Fasciclin II (DSHB #1D4, 1:100), mouse anti-βgal (DSHB #40-1a, 1:150), mouse anti-HA (Covance #MMS-101P-500, 1:1000), rabbit anti-HA (Covance #PRB-101C-500; 1:2000), Alexa Fluor 647 goat anti-rabbit (Jackson #123-605-021; 1:500), Cy3-conjugated goat anti-mouse (Jackson #115-165-003, 1:1000). Embryos were genotyped using balancer chromosomes carrying *lacZ* markers (CyO,wg), or by the presence of epitope-tagged transgenes. Nerve cords from embryos of the desired genotype and developmental stage were dissected and mounted in 70% glycerol/PBS. Fluorescent confocal stacks were collected using a Leica SP5 confocal microscope and processed by Fiji/ImageJ [61] and Adobe Photoshop software. For quantification of commissural clearance of Robo1 variants (Fig. 2), anti-HA pixel intensities of longitudinal and commissural axons were measured using Fiji/ImageJ and the ratio of commisural:longitudinal HA levels (averaged from 3 embryos per genotype) is reported as “Commissure HA levels” in Fig. 2. For quantification of Comm-dependent HA-Robo1 downregulation (Fig. 5), anti-HA pixel intensities of longitudinal axons were measured using Fiji/ImageJ and average HA intensities from embryos overexpressing Comm were compared to HA intensities from sibling embryos lacking Comm overexpression. The ratios of HA levels in UAS-Comm-positive embryos to HA levels in UAS-Comm-negative embryos is reported as “Comm-dependent HA downregulation” in Fig. 5.

## Acknowledgements

Stocks obtained from the Bloomington Drosophila Stock Center [National Institutes of Health (NIH) grant P40 OD-018537) and cultured cells obtained from the Drosophila Genomics Resource Center (NIH 2P40OD010949) were used in this study. Monoclonal antibodies were obtained from the Developmental Studies Hybridoma Bank, created by the Eunice Kennedy Shriver National Institute of Child Health and Human Development of the NIH and maintained at The Department of Biology, University of Iowa, Iowa City, IA 52242.

